# An evolutionarily conserved strategy for ribosome binding and inhibition by β-coronavirus non-structural protein 1

**DOI:** 10.1101/2023.06.07.544141

**Authors:** Stephanie F. Maurina, John P. O’Sullivan, Geetika Sharma, Daniel C. Pineda Rodriguez, Andrea MacFadden, Francesca Cendali, Morkos A. Henen, Jeffrey S. Kieft, Anum Glasgow, Anna-Lena Steckelberg

## Abstract

An important pathogenicity factor of SARS-CoV-2 and related coronaviruses is Nsp1, which suppresses host gene expression and stunts antiviral signaling. SARS-CoV-2 Nsp1 binds the ribosome to inhibit translation through mRNA displacement and induces degradation of host mRNAs through an unknown mechanism. Here we show that Nsp1-dependent host shutoff is conserved in diverse coronaviruses, but only Nsp1 from β-CoV inhibits translation through ribosome binding. The C-terminal domain of all β-CoV Nsp1s confers high-affinity ribosome-binding despite low sequence conservation. Modeling of interactions of four Nsp1s to the ribosome identified few absolutely conserved amino acids that, together with an overall conservation in surface charge, form the β-CoV Nsp1 ribosome-binding domain. Contrary to previous models, the Nsp1 ribosome-binding domain is an inefficient translation inhibitor. Instead, the Nsp1-CTD likely functions by recruiting Nsp1’s N-terminal “effector” domain. Finally, we show that a viral *cis*-acting RNA element has co-evolved to fine-tune SARS-CoV-2 Nsp1 function, but does not provide similar protection against Nsp1 from related viruses. Together, our work provides new insight into the diversity and conservation of ribosome-dependent host-shutoff functions of Nsp1, knowledge that could aide future efforts in pharmacological targeting of Nsp1 from SARS-CoV-2, but also related human-pathogenic β-coronaviruses. Our study also exemplifies how comparing highly divergent Nsp1 variants can help to dissect the different modalities of this multi-functional viral protein.

## Introduction

As obligate intracellular parasites, all viruses require the cellular gene expression machinery for replication, and many have evolved to be master manipulators of ribosomes and translation factors^1^. Viral manipulation of cellular gene expression serves multiple purposes during infection: *(1)* To divert cellular resources towards the preferential synthesis of viral proteins; *(2)* to maintain viral protein synthesis when activation of stress signaling pathways inhibits canonical translation initiation; and *(3)* to prevent the synthesis of antiviral proteins and peptides. The global inhibition of host gene expression during viral infection is commonly referred to as “host shutoff”. SARS-CoV-2, the causative agent of a life-threatening respiratory disease and the COVID-19 pandemic, and the closely related SARS-CoV use very efficient and multifaceted host shutoff strategies to rewire gene expression and inhibit antiviral signaling. The resultant inhibition of interferon (IFN) signaling is believed to contribute significantly to COVID-19 pathogenicity^2, 3^.

The best studied coronaviral host shutoff factor is the viral non-structural protein 1 (Nsp1), a ∼20 kDa N-terminal cleavage product of the coronavirus replicase polyprotein^4^. Foundational work on SARS-CoV revealed that Nsp1 inhibits host protein synthesis by preventing translation initiation and inducing the widespread cleavage and degradation of host mRNAs^5–9^. Both functions require an interaction of Nsp1 with the small ribosomal subunit^9^. Expression of Nsp1 in human cell culture recapitulates the translation shutdown and mRNA degradation phenotype, suggesting that no additional viral proteins are involved^9–11^. Recent single particle cryo-EM and single molecule biophysical studies revealed that a short C-terminal α-helical domain of SARS-CoV-2 Nsp1 binds to the mRNA entry channel of the mammalian 40S ribosomal subunit, thereby displacing the mRNA and inhibiting translation initiation^12–16^. Coronaviral mRNAs partially evade the Nsp1-dependent host shutoff, a function attributed to structured RNA elements in the viral 5’ UTR that function by an unknown mechanism ^5, 17–23^. Nsp1 also disrupts the nucleocytoplasmic export of mRNAs, likely through interactions with the mRNA export factor NXF1 and the nuclear pore protein Nup93, and causes a deregulation of stress granule biosynthesis^11, 24–28^. Through its multiple roles during infection, Nsp1 functions as an important virulence factor, exemplified by the fact that Nsp1-deletion mutants of SARS-CoV are highly attenuated in mice^29, 30^; consequently, Nsp1 has been explored as a target for antiviral strategies ^22, 31–36^. While a heightened interest in SARS-CoV-2 biology due to the ongoing COVID-19 pandemic has greatly accelerated the research^10, 11, 37, 38^, there are still many open questions about the underlying molecular mechanisms of Nsp1 functions.

Most research efforts to date have focused on Nsp1 from the related β-CoVs SARS-CoV and SARS-CoV-2, which share over 86% sequence identity and highly conserved functionality (Fig. 1A-B). We do not know if Nsp1-dependent host shutoff mechanisms are conserved in other coronaviruses. The *Coronaviridae* family is divided into four genera: α-, β-, ψ- and ο-coronaviruses, of which both α- and β-CoVs encode Nsp1. Incidentally, these are the genera that contain the seven human-pathogenic coronaviruses (the α-CoVs HuCoV-229E and HCo-NL63, and β-CoVs SARS-CoV, SARS-CoV2, MERS-CoV, HuCoV-HKU1 and HuCoV-OC43) (Fig. 1A). Surprisingly, Nsp1 is one of the least conserved non-structural proteins in the coronavirus genome, with only ∼20% sequence identity between most β-CoVs, and even less sequence conservation between the different genera, precluding automatic sequence alignments (Fig. 1B, Fig S1A). Nonetheless, three-dimensional structures - solved by x-ray crystallography and NMR - of several α- and β-CoV Nsp1 show a similarly folded N-terminal globular core, formed by a six-stranded β-barrel fold in the middle of two α-helices (Suppl Fig. 1B-D)^39–43^. In all β-CoV Nsp1s, this globular core is followed by a C-terminal tail, which includes the ribosome binding domain in SARS-CoV-2 Nsp1 ^12–15^. Several α- and β-CoV Nsp1s have been implicated in host shutoff^26, 39–41, 44–52^, but it is currently unknown if the ability to interact with and obstruct the ribosomal mRNA entry channel is conserved. Of note, MERS-CoV Nsp1, porcine epidemic diarrhea virus (PEDV) and transmissible gastroenteritis virus (TGEV) Nsp1 have all been shown to inhibit mRNA translation without interacting with the ribosome^44, 49, 50^, while Nsp1s from the α-CoV HuCoV-229E and HuCoV- NL63 were shown to interact with the ribosomal protein RPS6, despite missing a C-terminal domain^51^. These findings highlight mechanistic variability of Nsp1, and prompted us to investigate the ability of diverse Nsp1s to interact with and manipulate the cellular translation machinery.

**Figure 1:**
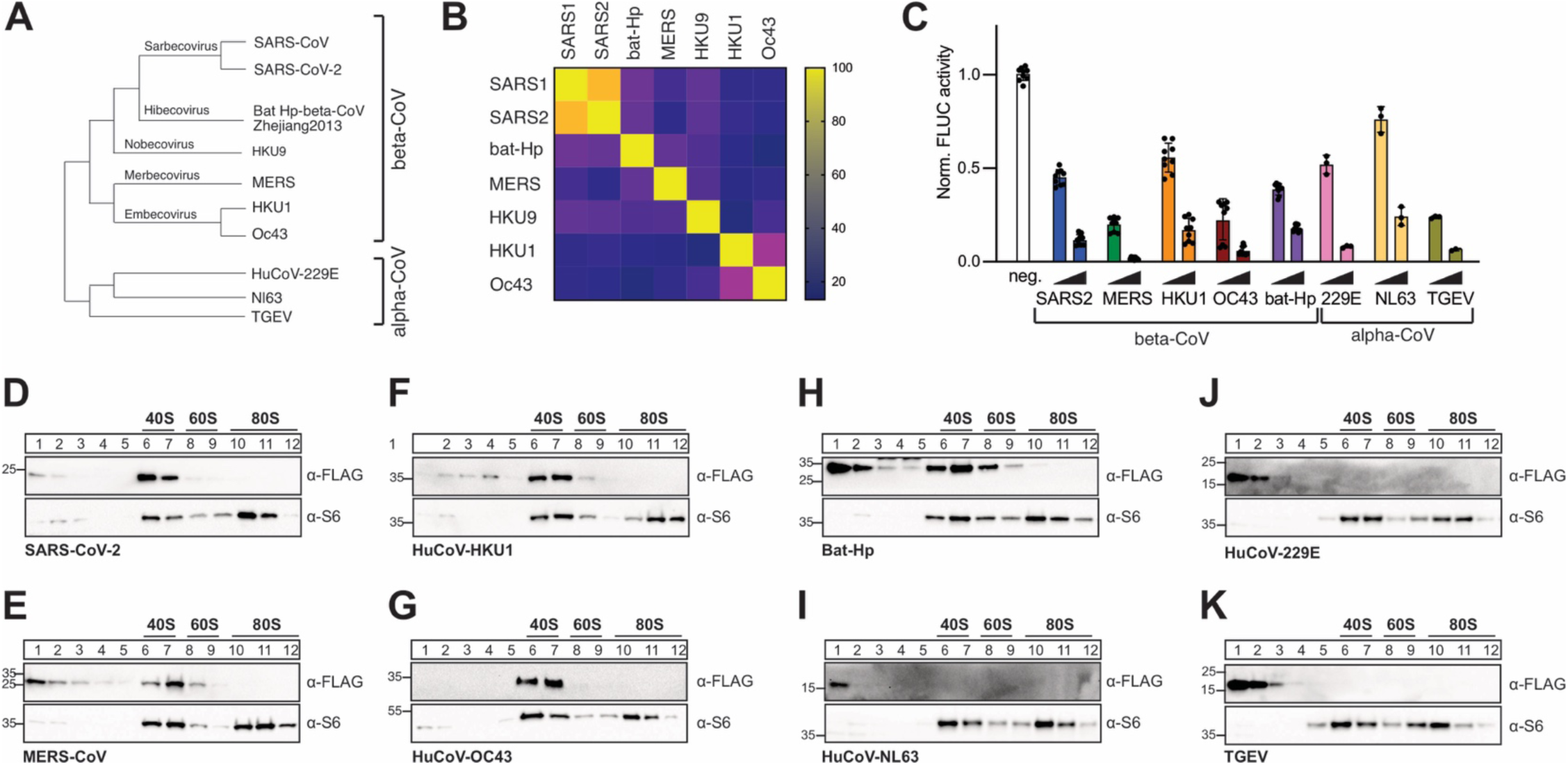
Conserved host shutoff and ribosome binding of coronaviral Nsp1. A) Evolutionary relationship of select α- and β-coronaviruses. The phylogenetic tree is based on RdRP conservation. B) Heat map of sequence variation in β-CoV Nsp1 proteins. C) Host shutoff in HEK293T cells transiently expressing Nsp1 from the indicated viruses, measured by luciferase activity of a co-expressed FLuc reporter gene. D-K) Polyribosome gradient analysis of HEK293T cell lysates expressing 3x-FLAG tagged Nsp1 from SARS-CoV-2 (D), MERS-CoV (E), HuCoV-HKU1 (F), HuCoV-OC43 (G), Bat Hp β-CoV Zhejiang2013 (H), HuCoV-NL63 (I), HuCoV-229E (J) or TGEV (K). Western blot analysis for the presence of Nsp1 (α-FLAG) and ribosomal protein RPS6 (α-S6).

We show that all tested proteins efficiently inhibit reporter gene expression in human cultured cells and mammalian cell extracts, but only Nsp1 from β-CoVs interact with the ribosome. The C- terminal domain (CTD) of β-CoV Nsp1 is necessary and sufficient to mediate this high-affinity ribosome binding. Using computational modeling, we show that the CTD from diverse viral proteins likely folds into very similar α-helical structures when complexed with the ribosome. Surprisingly, there are few absolutely conserved residues in the CTD, and conservation of surface charge and hydrophobic interactions, rather than amino acid sequence, maintains the high-affinity ribosome interaction. The specificity of host shutdown is likely achieved through additional functionality contained within the N-terminal protein domain. This model is supported by our observation that the C-terminal domain alone confers high-affinity ribosome binding, but only the full-length protein induces a strong host shutoff effect. In addition, we show that RNA structure elements of the SARS-CoV-2 5’ leader sequence regulate SARS-CoV-2 Nsp1 function, but have little or no effect on Nsp1 from other β-CoVs, indicating that a cis-acting viral RNA element has co-evolved with its cognate protein partner.

Taken together, our study provides evidence for an evolutionarily conserved mechanism by which Nsp1 from diverse β-CoVs binds to and inhibits the mammalian ribosome. The functional conservation of Nsp1 suggests that pharmacological targeting of the Nsp1-ribosome interaction could be a viable strategy not only against SARS-CoV-2, but also to safeguard against future outbreaks of related, human-pathogenic β-coronaviruses. Moreover, our study exemplifies how comparing highly divergent Nsp1 variants can help to dissect the different modalities of this multi-functional viral protein.

## Results and Discussion

### Nsp1-dependent host shutoff is conserved in α- and β-CoVs, but only β-CoV Nsp1s interact with ribosomes

We picked representative Nsp1 variants from four β-CoV subgenera and several highly divergent α-CoVs for a total of 8 Nsp1. These included the 4 common cold human coronaviruses HuCoV-HKU1, HuCoV-OC43 (both β-CoV, lineage A), HuCoV-229E and HuCoV-NL63 (both α-CoV), the 2 highly pathogenic human coronaviruses SARS-CoV-2 (β-CoV, lineage B), and MERS-CoV (β-CoV, lineage C), as well as Bat-Hp β-CoV Zhejiang2013 (β-CoV, the only known virus in the hibecovirus subgenus), and the pig-infecting model α-CoV TGEV (Fig. 1A). 3x-FLAG-tagged Nsp1 variants were transiently expressed in HEK293T cells, and host shutoff was monitored by measuring the activity of a co-transfected Firefly luciferase (FLuc) reporter. Compared to a vector control, all Nsp1 variants strongly inhibited FLuc expression in a concentration-dependent manner (Fig. 1C). The host shutoff effect was further confirmed by co-expressing a GFP reporter construct (Suppl. Fig. 1E). While all tested Nsp1s induced host shutoff, the efficiency varied considerably, with SARS-CoV-2, MERS-CoV, and bat-Hp Nsp1 consistently showing the strongest effects (Suppl. Fig. 1E).

We next investigated the ability of the same Nsp1 variants to interact with human ribosomes. Nsp1 was transiently expressed in HEK293T cells, cleared cell lysates separated through a sucrose gradient, and fractions containing ribosomal particles were analyzed by Western blotting for the presence of Nsp1 (Fig. 1D-K). We observed a striking dichotomy of ribosome interactions, with all β- CoV Nsp1s co-migrating with the 40S ribosomal subunits, and all of the tested α-CoV Nsp1s in the unbound fractions. These results contrast with previous studies showing that the β-CoV MERS-CoV Nsp1 does not interact with ribosomes^44^, and that Nsp1 from α-CoV HuCoV-229E and HuCoV-NL63 interact with the ribosomal protein RPS6^51^. The discrepancies might be due to differences in sample preparation and experimental procedure. A previous MERS-CoV Nsp1 ribosome interaction study was performed in the presence of detergent (1% Triton X-100), which might weaken Nsp1-ribosome interactions. When revisiting the previous MERS-CoV Nsp1 study, we furthermore observed faint MERS-CoV Nsp1 bands in fractions containing 40S ribosomal particles, but they were weaker than the corresponding bands from SARS-CoV Nsp1 and likely interpreted as “bleed-through” from the unbound fractions^44^. The previously described ribosome interactions of α-CoV HuCoV-229E and HuCoV-NL63 Nsp1s were based on co-immunoprecipitation of the ribosomal protein RPS6 with transiently expressed viral Nsp1. Co-immunoprecipitation assays are prone to nonspecific interactions and nonspecific antibody staining, and do not distinguish between interactions with individual proteins and the fully assembled ribosomal particle. In contrast, several independent studies confirmed that Nsp1 from the α-CoVs TGEV and PEDV did not interact with RPS6^49, 50^. Overall, we conclude that Nsp1s from diverse β-CoVs maintain the ability to interact with the small ribosomal subunit despite considerable sequence variation, while this functionality is not present in the significantly smaller (∼10 kDa) Nsp1 from α-CoVs. Nsp1 thus evolved to manipulate gene expression through both ribosome- dependent and -independent mechanisms.

### Ribosome binding through the β-CoV Nsp1 C-terminal domain

The ability to interact with the small ribosomal subunit coincided with the presence of the Nsp1 C- terminal domain, which is also the protein domain bound to the mRNA entry channel in SARS-CoV-2 Nsp1-ribosome structures^12, 13^. To test if the C-terminus mediates ribosome binding in all β-CoV Nsp1s, C-terminally truncated mutants of SARS-CoV-2, MERS-CoV, HuCoV-HKU1, HuCoV-OC43 and bat-Hp Nsp1 were expressed in HEK293T cells and polysome profiles resolved as described above (Fig. 2B). None of the C-terminally truncated Nsp1 mutants co-migrated with ribosomal particles, indicating that the C-terminal domain is indeed necessary for ribosome binding. Furthermore, the Nsp1 CTD alone was sufficient to confer ribosome binding when C-terminally fused to GFP, as shown by 40S co-migration of GFP fused to the C-terminal domain of any β-CoV Nsp1, but not of GFP alone (Fig. 2C). Of note, the CTD of MERS-CoV Nsp1 showed the weakest ability to shift GFP to the 40S fractions, suggesting that its binding might be weaker than that of other Nsp1 variants. Taken together, the C-terminus of Nsp1 from highly diverse β-CoVs confer binding to the human small ribosomal subunit.

**Figure 2:**
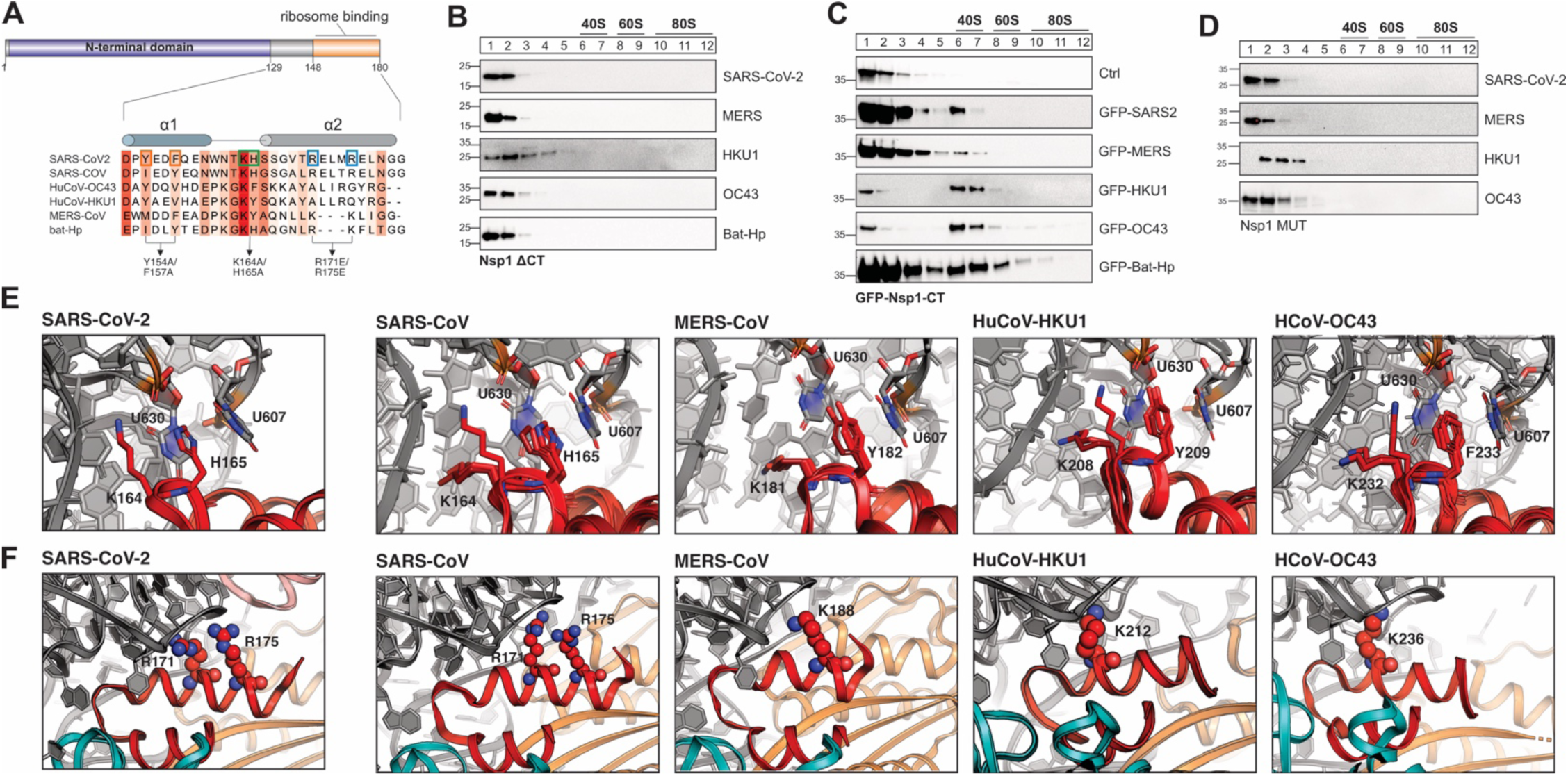
A conserved α-helical structure in the C-terminus of β-CoV Nsp1 mediates ribosome binding. A) Sequence alignment of the C-terminal domain of Nsp1 from SARS-CoV, SARS-CoV-2, MERS-CoV, HuCoV-HKU1, HuCoV-OC43 and bat-Hp Nsp1. The Sequence conservation is indicated by red shading. Previously characterized mutants in the C-terminus of SARS-CoV-2 Nsp1 are indicated by boxes and arrows. B-D) Polyribosome gradient analysis of HEK293T cells transiently expressing the indicated 3x-FLAG-tagged Nsp1 variants: C-terminally truncated Nsp1 mutants (B), GFP (top) or GFP fused to the CTD of Nsp1 from the indicated viruses (C), and Nsp1 variants in which the conserved KH or KY/F motif was mutated (D). Western blot analysis: α-FLAG. E) Details of the interactions between the KH-motif of the SARS-CoV-2 Nsp1 CTD and the ribosome (left, pdb 6zlw), and modeled interactions between the KH-motif of SARS-CoV (2^nd^ panel), KY motif of MERS-CoV (3^rd^ panel) and HuCoV-HKU1 (4^th^ panel), and the KF-motif of HuCoV-OC43 Nsp1 (right) with the 18S rRNA. The aromatic residues of the KY/F-motif likely form stacking interactions with ribosomal U607 and U630, similar H165 in SARS-CoV-2 Nsp1. Nsp1 in red and 18S rRNA in grey. F) Details of the interactions between helix α2 of SARS-CoV-2 Nsp1 and the 18S rRNA of the small ribosomal subunit (left, pdb 6zlw). Details of the modeled interactions between helix α2 of Nsp1 from the indicated viruses and the 18S rRNA (right). Note that the electrostatic interactions of R171 and R175 are replaced by positively charged amino acids at different positions along α2 in MERS-CoV, HuCoV-HKU1 and HuCoV-OC43 Nsp1. Nsp1 is shown in red, 18S rRNA in grey and ribosomal proteins uS3 in teal and uS5 in orange.

All early structures of SARS-CoV and SARS-CoV-2 Nsp1 are missing the CTD, suggesting that the C-terminus of Nsp1 is intrinsically disordered in solution^42, 43, 53, 54^, and adopts its ordered helix-loop-helix fold only when bound to the small ribosomal subunit^12, 13^. To gain further insight into the dynamics of full-length Nsp1 in solution, we measured R_1_ and R_2_ relaxation rates of full length Nsp1 from SARS-CoV-2 by NMR, to study dynamics in a ps to µs time scale. Analysis of R_2_ relaxation data showed an expected decrease in relaxation rates after amino acid position G-129, demonstrating that the entire CTD is intrinsically disordered in solution. Closer inspection of the dynamic C-terminus revealed several residues (His-134, Ser-141, Asp-152 and Thr-170) with above average R_2_ rates, indicating potential structural compaction through stabilizing intramolecular interactions of these residues. These data are in agreement with other NMR studies on full-length SARS-CoV-2 Nsp1, published while our manuscript was in preparation^55, 56^.

### Molecular variability of Nsp1’s ribosome-binding domain

The observation that a short (<30 amino acids), intrinsically disordered and poorly conserved C-terminal tail of Nsp1 is sufficient for ribosome binding was surprising, especially in the context of current mechanistic models suggesting that ribosome binding promotes efficient translation shutdown through obstruction of the mRNA entry channel^16^. While the Nsp1-CTD is intrinsically disordered in solution, recent cryo-EM studies revealed that when complexed with mammalian 40S ribosomal subunits, the most C-terminal residues of SARS-CoV-2 Nsp1 (residues 148-180) forms two short α-helices that bind to the ribosomal mRNA entry channel^12–15^. To get a clearer picture of the ribosome interactions of the CTD from other Nsp1 variants, we performed homology modeling of Nsp1 from SARS-CoV, MERS- CoV, HuCoV-HKU1 and HuCoV-OC43 bound to the human small ribosomal subunit using the Rosetta software suite^57^ and eight SARS-CoV-2 Nsp1 structures^12^ as templates (Suppl. Figure 3A). As expected, Nsp1 from the closely related SARS-CoV was predicted to form a 2-helix structure, but so were the C-terminal domains from the more divergent viruses MERS-CoV, HuCoV-HKU1 and HuCoV-OC43 (Suppl. Fig 3B). The length of α2 varies slightly, with α2 of SARS-CoV and SARS-CoV-2 being the longest (13 residues), and the short α2 of the MERS-CoV model being and a full helical turn shorter (9 residues). In all cases, the two helices stabilize each other against the ribosomal protein uS5 through hydrophobic residues at the helix interface (Suppl. Fig. 3C). The loop contains the conserved KH motif in SARS-CoV and SARS-CoV-2 Nsp1 – key residues that form salt bridges and stacking interactions with the h18 loop in the 18S rRNA^12, 13^. The same position has a KY motif in MERS-CoV and HuCoV-HKU1, and a KF motif in HuCoV-OC43 Nsp1 (Fig. 2A, Suppl. Fig. 3B); these residues can likely partially substitute for the rRNA interactions of KH in SARS-CoV and SARS-CoV-2.

Analysis of the electrostatic potential on the surface of the SARS-CoV-2 Nsp1 CTD has revealed 3 distinct patches that collectively mediate high-affinity binding to the ribosomal mRNA entry channel: (1) a negatively charged patch on α1, which interacts with positively charged residues of ribosomal protein uS3, (2) a hydrophobic patch on the α1/α2 interface which interacts with hydrophobic side chains on ribosomal protein uS5, and (3) a positive patch on α2 which interacts with rRNA^12^. The electrostatic surface potential of SARS-CoV-2 Nsp1-CTD is partially maintained in the models of SARS- CoV, MERS-CoV, HuCoV-OC43 and HuCoV-HKU1 Nsp1-CTD (Fig. 2E-F), suggesting that a conservation of surface charge, rather than a specific amino acid sequence, might primarily contribute to ribosome binding. The importance of the conserved hydrophobic surface of the α1/α2 interface facing uS5 (Fig. S3E) is corroborated by the fact that mutating hydrophobic residues in α1 of SARS-CoV-2 Nsp1-CTD (Y154A/F157A) completely abolished ribosome binding^13^.

The most important interactions between the Nsp1 CTD and the small ribosomal subunit involve a positive surface patch on the α2 helix (Suppl. Fig. 3F). In SARS-CoV-2, multiple positively charged amino acids in the CTD of Nsp1 tightly bind to h18 of the 18S rRNA: In particular, K164 interacts with the rRNA phosphate backbone, and H165 stacks between U607 and U630, in addition to forming electrostatic interactions with the phosphate backbone of the rRNA^12, 13^ In MERS-CoV, HCoV-HKU1 and HuCoV-OC43 Nsp1-CTD, H165 in the loop is replaced by a less polar aromatic amino acid (Tyr or Phe). Our modeling suggests that the KY/F motif likely interacts with the 18S rRNA in a very similar manner to the KH-motif in SARS-CoV-2 Nsp1, with the lysine forming electrostatic interactions with the rRNA phosphate backbone, and the aromatic residue stacking between U607 and U630 (Fig. 2E). In line with a conserved role for the KH- and KY/F motif, replacing the 2 residues with alanine in Nsp1 from SARS-CoV-2, MERS-CoV, HuCoV-HKU1 and HuCoV-OC43 completely abrogated ribosome binding in human cultured cells, as shown by polyribosome gradient analysis (Fig. 2D).

In addition to the KH-motif, interactions between the 18S rRNA and R171 and R173 from SARS-CoV-2 are crucial for ribosome binding^13^. Surprisingly, these residues are not conserved in the more divergent Nsp1 variants we studied (Fig. 2A). In HuCoV-HKU1 and HuCoV-OC43 Nsp1 models, the loss of R171 and R173 appears to be compensated by lysine residues in the α2 helix. Interestingly, these lysine residues are positioned in the first helical turn of α2, whereas R171 and R173 are in the second and third helical turn in α2 from SARS-CoV-2, respectively. Consequently, the positively charged residues in α2 are poised to interact with slightly different sections of 18S rRNA backbone (Fig. 2F). In MERS-CoV, a two adjoining arginine residues in the second helical turn of α2 leads to a weaker overall positive surface charge and likely a weaker interaction with the 18S rRNA (Fig. 2F, Suppl. Fig. 3F). Finally, bat-Hp Nsp1 appears to use a mixed strategy to interact with 18S rRNA: It has a shorter α2 helix, similar to MERS-CoV Nsp1, but a KH-motif in the loop, like SARS-CoV and SARS-CoV-2; the KH-motif can potentially anchor the C-terminal domain by forming additional electrostatic interactions with the phosphate backbone of the RNA. Importantly, our homology modeling data is corroborated by a contemporaneous preprint describing the cryo-EM structures of ribosome-bound MERS-CoV and bat-Hp Nsp1^58^.

Collectively, our study highlights remarkable variability in the molecular interactions between the C-termini of β-CoV Nsp1s and the ribosomal mRNA entry channel. The ribosomal mRNA entry channel is highly conserved from yeast to humans (including amino acids R116, R117 and R143 of ribosomal protein uS3, and nucleotides U607 and U630 of the 18S rRNA, which form direct interactions with the SARS-CoV-2 Nsp1 CTD^12, 13^). Relatively unspecific electrostatic interactions with a conserved ribosomal binding pocket likely allow Nsp1 to tolerate considerable sequence variability in the CTD without losing its host shutoff capacity. Consistent with a pivotal role of the ribosomal mRNA entry channel for Nsp1 binding, a recent study showed that mutating a key residue on the Nsp1-interacting interface of uS3 (R116D) rendered ribosomes immune to Nsp1-dependent translation inhibition^59^.

### The Nsp1-CTD confers high-affinity binding but is an inefficient translation inhibitor

Previous studies have proposed that Nsp1 inhibits translation initiation by competing with mRNA for binding to the mRNA entry channel^12, 13, 16^. To correlate ribosome binding to translation shutdown, we tested the ability of the C-terminal domain of several viral Nsp1s fused to GFP to inhibit FLuc reporter gene expression in human cell culture (Fig. 3A-B). All GFP-CTD fusion proteins reduced luciferase reporter activity, but they were significantly less efficient in inhibiting gene expression than the full length versions of each protein. This suggests that ribosome binding alone is inefficient in promoting translation inhibition, and that additional functionality contained in the N-terminal domain is essential to achieve the full host shutoff effect. In line with this hypothesis, previous studies have identified conserved residues in the N-terminal domain (NTD) of SARS-CoV and SARS-CoV-2 that are required to induce the degradation of ribosome-associated host mRNAs^9–11, 37^.

**Figure 3:**
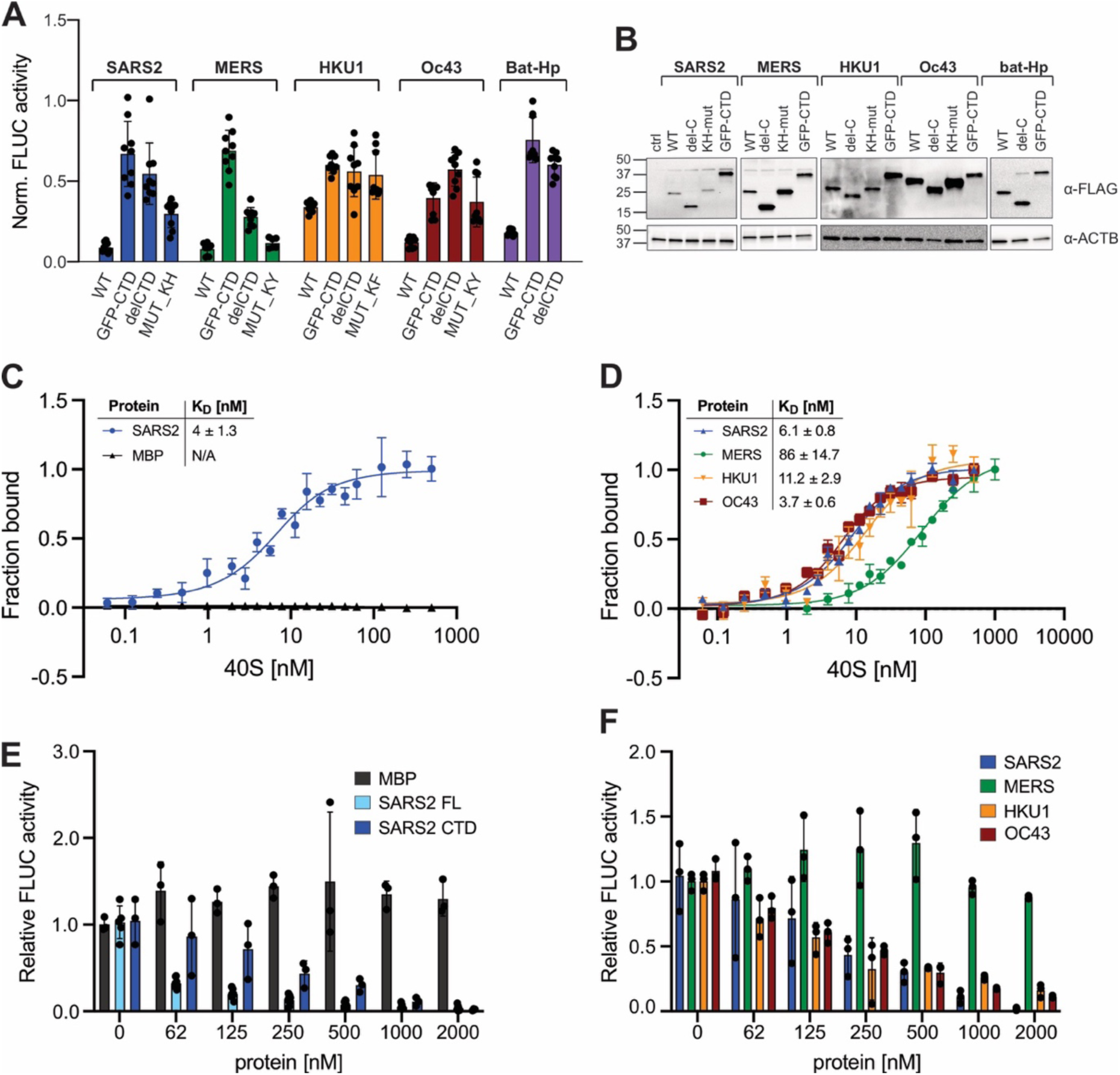
Correlation between ribosome binding and translation inhibition in different β-CoV Nsp1s. A) Host shutoff in HEK293T cells transiently expressing 3xFLAG-tagged wild type or mutant Nsp1 from the indicated viruses, measured by FLuc activity. B) α-FLAG immunoblot analysis of HEK293T cells from (A). C) Equilibrium binding measurements of FAM-labeled SARS-CoV-2 Nsp1 to purified rabbit 40S ribosomal subunits, maltose binding protein (MBP) served as a negative control. n=3, error bars=SEM. D) Equilibrium binding measurements of FAM-labeled MBP fused to the Nsp1-CTD from the indicated viruses to rabbit 40S subunits. n=3, error bars=SEM. E) Cell free translation of FLuc reporter mRNA in rabbit reticulocyte lysate supplemented with increasing concentrations of MBP, SARS-CoV-2 Nsp1 (“SARS2 FL”), or MBP fused to the CTD of SARS-CoV-2 (“SARS2 CTD”). n=3, error bars=SEM. F) Cell free translation of an FLuc reporter mRNA in the presence of increasing concentrations of MBP fused to the Nsp1 CTD of the indicated viruses. n=3, error bars=SD.

We next tested the ability of ribosome-binding deficient Nsp1 mutants (containing either deletions of the entire C-terminal domain, or mutants of the KH- or KY/F-motif) to inhibit FLuc reporter gene expression. In the case of SARS-CoV-2, HuCoV-HKU1, HUCoV-OC43 and bat-Hp Nsp1, abrogating ribosome binding severely reduced, but did not completely abolish, the host shutoff effect of Nsp1 (Fig. 3A-B). This suggests that Nsp1 from these viruses heavily relies on ribosome binding, but can additionally inhibit gene expression in a ribosome-independent manner. Possible mechanisms for ribosome-independent host shutoff are the previously described roles of Nsp1 in mRNA export and cytoplasmic granule formation^11, 24–28^. Of note, mutating the ribosome-binding domain of MERS-CoV Nsp1 only moderately reduced its host shutoff function, suggesting that even though we observed ribosome binding of MERS-CoV Nsp1 in human cells, this interaction is dispensable for translation inhibition. Our observation corroborates earlier studies describing that MERS-CoV Nsp1 can inhibit protein synthesis without binding to the ribosome^44^. Future studies are needed to reveal how the NTD of Nsp1 contributes to host shutoff, and how Nsp1 proteins (especially MERS-CoV Nsp1) can inhibit gene expression in ribosome-independent manners. Of note, a recent study comparing the transcriptome of cells expressing Nsp1 from SARS-CoV, SARS-CoV-2, MERS-CoV and HuCoV-229E discovered the largest overlap between MERS-CoV and HuCoV-229E Nsp1 – potentially indicating similar targeting mechanisms of proteins that do not require ribosome-binding for function^60^.

Direct comparison of the host shutoff effect of Nsp1 mutants is complicated, because expression levels of different Nsp1 constructs in HEK293T cells varied considerably (Fig. 3B). To reliably relate translation shutoff to ribosome binding affinity of Nsp1, we expressed and affinity-purified recombinant full-lengths SARS-CoV-2 Nsp1 from *E. coli.* Maltose binding protein (MBP) was purified similarly to serve as a negative control. Since we had previously shown that the CTDs of Nsp1 from SARS-CoV- 2, MERS-CoV, HuCoV-HKU1 and HuCoV-OC43 are both necessary and sufficient to promote ribosome binding, and that the Nsp1 CTD can confer ribosome binding when fused to a different protein (GFP), we also purified MBP fused to the C-terminal domains of Nsp1 from SARS-CoV-2, MERS-CoV, HuCoV-HKU1 and HuCoV-OC43. All proteins were site-specifically fluorescein-labeled and binding to small (40S) ribosomal subunits purified from rabbit reticulocyte lysate was measured through fluorescence anisotropy. (Fig. 3C-D).

Full length SARS-CoV-2 Nsp1 bound tightly to 40S subunits (dissociation constant (K_D_) of ∼4 nM), similar to binding affinities previously reported from single-molecule experiments^16^, and significantly lower than the equilibrium binding affinity of SARS-CoV-2 Nsp1 with 80S ribosomes (K_D_∼500 nM)^10^. We did not observe the binding of maltose binding protein (MBP) to purified 40S subunits, confirming that the binding is specific. Interestingly, MBP fused to the CTD of SARS-CoV-2 Nsp1 (MBP- CTD_SARS2_) HuCoV-HKU1 (MBP-CTD_HKU1_) and HuCoV-OC43 (MBP-CTD_OC43_) showed similar 40S binding affinities as full-length SARS-CoV-2 Nsp1, demonstrating that C-terminal domain alone is sufficient to convey high affinity ribosome binding and that the ability to bind tightly to the mRNA entry channel is maintained in all proteins, despite considerable molecular variability of the ribosome-binding domains (Fig. 3D). It should be noted that limiting protein concentrations in our equilibrium binding assays were near the apparent K_D._ We fitted binding to a quadratic equation to account for an intermediate binding regiment^61^, but it is possible that the reported K_D_ values underestimate the real affinities. The only protein that displayed considerably weaker, albeit still remarkably strong, 40S binding was MBP-CTD_MERS_ (K_D_ ∼ 86 nM). This is consistent with our cell-based assays (Fig. 2C). In accordance with electrostatic interactions being a major contributor to Nsp1-40S binding, we observed significantly higher dissociation constants upon increasing the concentration of K^+^ in the buffer from 120 mM to 250 mM (∼30 nM, Suppl. Fig. S3I).

We next analyzed the ability of the recombinant proteins to inhibit reporter mRNA translation in a cell free translation system. To this end, FLuc mRNA was translated in rabbit reticulocyte lysate supplemented with increasing concentrations of the recombinant proteins. Despite having similar 40S- binding affinities, there was a striking difference in the ability of full-length Nsp1 and the MBP-CTD_SARS2_ to inhibit FLuc translation (Fig. 4E). Full length SARS-CoV-2 Nsp1 had a half-maximal inhibitory effect at <60 nM, whereas MBP-CTD_SARS2_ only reached a similar translation inhibition at ∼250 nM (Fig. 4E), further strengthening our hypothesis that important protein functionality of the Nsp1 NTD is required to achieve full translation shutdown strength. MBP alone had no effect on FLuc translation, confirming specificity of the effect. Importantly, our cell free translation assay is not confounded by translation- independent effects of Nsp1, such as mRNA export or mRNA granule formation; any observed differences in luciferase activity therefore directly relate to the ability of each protein to regulate mRNA translation or mRNA stability. We propose that the translation inhibition of MBP-CTD_SARS-CoV-2_ exclusively derives from its ability to displace the mRNA from the ribosomal mRNA entry channel and directly correlates with its 40S binding affinity. This hypothesis is supported by the fact that MBP- CTD_OC43_ and MBP-CTD_HKU1_ have similar half-maximal translation inhibitory concentrations in cell free translation assays (Fig. 4F). In contrast, MBP-CTD_MERS_, which binds 40S subunits with significantly lower affinity, was unable to suppress reporter mRNA translation in rabbit reticulocyte lysate, even at the highest tested concentrations (2 µM) (Fig. 4F).

**Figure 4:**
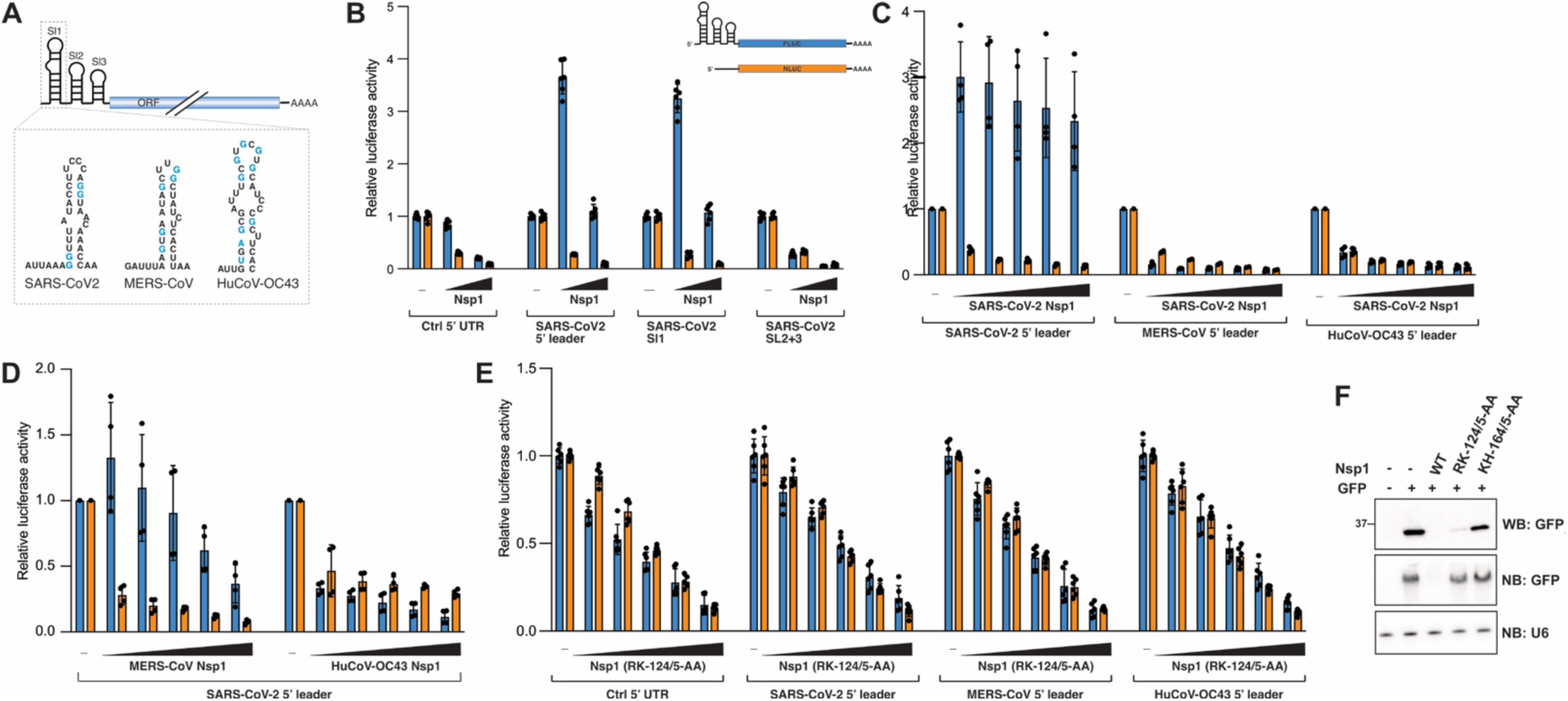
A virus-specific RNA-protein interaction protects viral RNA from Nsp1-dependent host shutoff. A) Scheme of the coronavirus 5’ leader sequence with proposed 2D structures of SL1 from SARS-CoV-2, MERS-CoV and HuCoV-OC43 shown below. SL1 from different β-CoVs shows remarkable sequence variability. B-E) Nsp1-dependent host shutoff or translation boost in HEK293T cells transiently expressing Nsp1 from the indicated viruses and FLuc reporters carrying the indicated 5’ UTRs. NLuc was co-transfected as an internal control. N=6, error bars = SD. F) Western blot of whole cell lysate (top, antibody: anti-GFP), and Northern blot of total RNA from HEK293T cells transiently expressing SARS-CoV-2 Nsp1 wild type or the indicated mutants and a GFP reporter. Probes are against GFP and U6 spliceosomal RNA.

Collectively, our experiments in human cell culture and *in vitro* show that the C-terminal domains of SARS-CoV-2, HuCoV-HKU1 and HuCoV-OC43 Nsp1 all bind to the small ribosomal subunit with equally high affinity despite significant sequence variation, and that this interaction alone is sufficient to inhibit translation at high protein concentrations. However, the translation shutdown is substantially more pronounced when the C-terminal ribosome binding domain is fused to the NTD of Nsp1. We thus propose an updated model for Nsp1 function, wherein the CTD’s primary role is not to inhibit translation, but to recruit Nsp1 (or more specifically the Nsp1 NTD) to its site of action. Consistent with the NTD as Nsp1’s main “effector” domain, multiple studies recently identified residues within the Nsp1-NTD that are crucial for host shutoff^7, 10, 62^. The molecular functions of these N-terminal residues are still poorly understood. Of note, a recent preprint shows that the NTD of bat-Hp Nsp1 binds to the decoding center of the 40S subunit, yet the same interactions were not observed for SARS-CoV-2 and MERS-CoV Nsp1^58^, while another preprint suggests that Nsp1 itself functions as an endonuclease to cleaves ribosome-associated mRNAs^63^.

Remarkably, despite the fact that MERS-CoV Nsp1 can bind ribosomes in cultured human cells, and that the CTD of MERS-CoV Nsp1 binds to purified ribosomal subunits *in vitro*, our investigation confirmed that MERS-CoV Nsp1-dependent host shutoff is less reliant ribosome binding^44^. It will be interesting to determine how MERS-CoV Nsp1 has lost the ribosome-dependency of host shutoff.

### A cis-acting protective viral RNA element has coevolved with SARS-CoV-2 Nsp1

Translation of coronaviral mRNAs continues in the presence of SARS-CoV, SARS-CoV-2 and MERS-CoV Nsp1, and an Nsp1-protective *cis*-acting RNA elements was identified in the viral 5’ leader sequence^5, 11, 15, 18–21, 23, 37, 44, 47^. The 5’ leader is an approximately 70-nucleotide long untranslated sequence appended to the 5’ end of all coronaviral mRNAs through a complex replication strategy known as discontinuous transcription^64^. It contains three small proposed stem loop structures (SL1, SL2 and SL3), of which SL1 was shown to be both necessary and sufficient to protect mRNAs from Nsp1-induced translation inhibition and degradation^18, 23, 47^. The ability to form SL1 is conserved across related β-CoVs, yet the sequence and structural similarities between viral leader sequences are low (Fig. 4A). Nevertheless, SARS-CoV SL1 can functionally replace its counterpart in the Mouse hepatitis virus (MHV) genome to produce viable chimeric viruses^65^, suggesting that SL1 might have similar functions in related viruses.

We sought to determine whether 5’ leader sequences from different β-CoVs function in a similar manner to the SARS-CoV-2 SL1 to regulate Nsp1 activity, and whether these sequence elements can provide cross-protection against Nsp1 from related viruses (Fig. 4). We replaced the 5’ UTR of our FLuc reporter construct with different viral sequences, and expressed the reporter constructs in HEK293T cells together with varying Nsp1 concentrations and a Nano-Luciferase (NLuc) reporter with a control 5’ UTR as an internal control. Consistent with previous reports^18, 19^, we observed that the SARS-CoV-2 5’ leader sequence not only confers protection from Nsp1, but in fact boosts reporter gene expression in an Nsp1-dependent manner, and that this functionality is contained in the SL1 sequence of the 5’ leader (Fig. 4B). Importantly, this translational boost appears to be specific to wild type Nsp1 and was not observed in the presence of an Nsp1 version with a mutated basic surface patch, which is also unable to induce mRNA degradation (SARS2-Nsp1_RK-124/5-AA) (Fig. 4E-F)^7, 10, 18^. Interestingly, neither the 5’ leader sequence from MERS-CoV, nor that of HuCoV-OC43, could provide Nsp1 cross- protection or an Nsp1-dependent translation boost when co-transfected with Nsp1 from SARS-CoV-2. Similarly, the SARS-CoV-2 5’ leader sequence provided a weaker cross-protection against Nsp1 from MERS-CoV, and no protection against Nsp1 from HuCoV-OC43 (Fig. 4C-D).

Collectively, these data suggest that RNA elements in the viral 5’ UTR have co-evolved to fine-tune the activity of their cognate viral protein, but have little to no effect in conjunction with Nsp1 from divergent β-CoVs. This observation is consistent with a recent study showing that even between the closely related sarbecoviruses SARS-CoV and SARS-CoV-2 – which have highly conserved Nsp1 and SL1 sequences -stronger interactions are observed between Nsp1 and its cognate viral RNA, an effect that can be partially rescued by mutating viral protein or viral RNA to resemble their cognate counterpart^47^.

The 5’ leader sequences from SARS-CoV-2, MERS-CoV and HuCoV-OC43 had an equally strong protective effect against MERS-CoV Nsp1 (Suppl. Fig. S4A), suggesting that some general structural feature of the 5’ UTR, rather than specific RNA-protein interactions, might regulate the MERSzCoV Nsp1-dependent translation shut-down. Analogous to SARS-CoV-2 Nsp1, the 5’ leader-dependent protection against MERS-CoV Nsp1 was only observed for wild type protein, but not for a mutant version, in which the highly conserved RK-motif was absent (MERS-Nsp1_RK-146/7-AA) (Suppl. Fig. S4C). Of note, an earlier study proposed that MERS-CoV Nsp1 selectively targets mRNA of nuclear origin, while sparing transcripts directly delivered to or synthesized in the cytoplasm (such as viral mRNAs)^44^. While we did not test cytoplasmic-derived mRNA in our study, it is possible that MERS-CoV Nsp1 employs multiple strategies to distinguish between cellular and viral transcripts.

In contrast, wild type HuCoV-OC43 Nsp1 was immune against all viral 5’ UTRs tested (Suppl. Fig. S4B), and it is currently unknown if and how HuCoV-OC43 ensures synthesis of viral proteins in the presence of Nsp1. It should be noted that HuCoV-OC43 was significantly less efficient in promoting host shutoff than Nsp1 from SARS-CoV, SARS-CoV-2 or MERS-CoV in all our reporter assays, it is therefore conceivable that HuCoV-OC43 can simply outcompete Nsp1 function by synthesizing large amounts of viral mRNAs.

Taken together, the RNA-protein interactions that fine-tune Nsp1 function appear specific to SARS-CoV-2 and the closely related SARS-CoV, but not present in more divergent β-CoVs. The uniqueness of this interaction is relevant for the design and interpretation of ongoing studies that explore therapeutic antisense oligos and small molecule inhibitors to target SARS-CoV-2 SL1 as antiviral strategies against COVID-19^18, 22^.

In agreement with several recent studies^19, 66^, we find that translation of reporter mRNAs carrying a viral 5’ UTR sequence is significantly increased in the presence of wild type Nsp1. The simplest explanation for the Nsp1-dependent translation boost is less competition for limiting translation factors upon Nsp1-dependent degradation of cellular transcripts. Alternatively, Nsp1 might function as a co-factor for non-canonical translation initiation of mRNAs carrying the viral 5’ leader sequence. In line with the latter hypothesis, a recent study showed that SARS-CoV-2 Nsp1 promotes cap-independent translation initiation^67^.

Our study provides new insight into the diversity and conservation of Nsp1-dependent host shutoff, strengthening the notion that antiviral therapies targeting Nsp1 could be a viable strategy against SARS-CoV-2 and related human- or animal-pathogenic coronaviruses. While many open questions remain, our comparative study exemplifies how the analysis of related viruses can identify conserved interactions and functions. We propose that analogous future studies will allow us to fill the remaining knowledge gaps about the coronaviral host shutoff factor Nsp1.

## Materials and Methods

### Plasmids

Constructs for protein expression in mammalian cell culture were inserted between NheI and NotI restriction sites of the pcDNA3.1 vector (Invitrogen). All Nsp1 expression constructs were cloned with an N-terminal 3x-FLAG tag. Firefly luciferase (FLuc) reporter constructs were cloned between NheI and NotI restriction sites in pcDNA3.1, and where indicated, the endogenous 5’ UTR sequence was replaced by viral sequences. Nano luciferase (NLuc) was expressed from pNL1.1.PGK (Promega). Constructs for protein expression in *E.coli* were cloned with an N-terminal 6xHis-tag into pET15b. Nsp1- delCTD constructs comprised residues 1-128 (SARS-CoV-2), 1-149 (MERS-CoV), 1-178 (HuCoV-HKU1), 1-196 (HuCoV-OC43), and 1-131 (bat-Hp). GFP and MBP were fused to 126-180 (SARS-CoV-2), 149-193 (MERS-CoV), 180-222 (HuCoV-HKU1), 196-246 (HuCoV-OC43) and 130-174 (bat-Hp).

### Cell culture and transfection

HEK293T cells were grown in grown in Dulbecco’s Modified Eagle Medium (DMEM), supplemented with 1X penicillin/streptomycin and 10% FBS at 37°C and 5% CO2. Cells were transfected using linear polyethylenimine (PEI) (25k, Polysciences, Inc) at a ratio of 2:1, and harvested 24 or 48 hours after transfection. Whenever cells were transfected with varying concentrations of pcDNA3.1-Nsp1, empty pcDNA3.1 vector was added to ensure equal DNA concentrations for all conditions. Cells for luciferase reporter assays were seeded in 48-well plates and transfected with 500 ug DNA plasmid (25 ng pcDNA3.1-FLuc, 25 ng pNL1.1.PGK, 50 ng pcDNA3.1-EGFP and between 25-400 ng pcDNA3.1-Nsp1 and up to 400 ng empty pcDNA3.1). Cells for sucrose gradient analysis were seeded in 15 cm plates (2 per gradient) and transfected with 15 ug pcDNA3.1-Nsp1 and 1.5 ug pcDNA3.1-EGFP. Cells for Northern blot analysis were seeded in 6-well plates and transfected with 1.5 ug pcDNA3.1-Nsp1 and 0.5 ug pcDNA3.1-EGFP.

### Sucrose gradient centrifugation

Transfected cells were lysed in 150 mM KOAc, 25 mM HEPES (pH 7.5), 5 mM Mg(OAc)_2_, 1% Triton-X 100, 2 mM DTT, 1x proteinase inhibitor at 24 hours after transfection. Lysates were cleared by centrifugation at 10,000xg for 10 min at 4C, and 2 mg cleared lysate was loaded onto a 10%-50% continuous sucrose gradient in 150 mM KOAc, 25 mM HEPES (pH 7.5), 5 mM Mg(OAc)_2_, 2 mM DTT and separated by ultracentrifugation at 36,000 rpm, 4 °C for 2:45 hrs in an SW 41 rotor. Fractions were collected using the BioComp Fractionator and analyzed by immunoblot analysis.

### Immunoblot analysis

Cells were lysed in 1x RIPA buffer (50 mM Tris (pH 7.5), 150 mM NaCl, 0.1% SDS, 0.5% sodium deoxycholate, 1% Triton X-100, 1x proteinase inhibitor cocktail (Sigma)). Protein concentrations were determined by BCA Assay (Pierce), and immunoblot analysis was performed using 20 μg whole cell lysate. After SDS-PAGE, proteins were transferred to nitrocellulose membranes blocked with EveryBlot Blocking Buffer (Bio-rad) and stained with specific antibodies diluted in TBS-Tween supplemented with 3% BSA. Specific antibodies were anti-DYKDDDDK (Invitrogen, #FG4R) anti-RPS6 (Invitrogen, #9HCLC), anti-β-actin (Abcam, #8226). For sucrose gradient experiments, 25 µl of each fraction was analyzed as described above.

### Luciferase assays

Wild-type and mutant Nsp1 proteins were overexpressed in HEK 293T cells, and luciferase reporter assays were performed using the Dual-Luciferase Reporter Assay System (Promega) according to manufacturer’s instructions and analyzed using a Biotek Synergy Neo2 multi-mode microplate reader.

### Protein expression and purification

All recombinant proteins were 6XHis-tagged, expressed in *E. coli* LOBSTR cells and purified using Ni-NTA resin (Thermo Fisher) followed by size exclusion chromatography on a Sephacryl S-100 column. The final product was aliquoted and stored at −80°C in buffer containing 20 mM Hepes-KOH pH 7.5, 120 mM KOAc, 5 mM Mg(OAc)_2_, 1 mM DTT. The purity of the recombinant proteins was verified by SDS-PAGE and Coomassie staining.

### FAM-labeling of recombinant proteins

All recombinantly expressed and purified proteins contained a single N-terminal cysteine residue to allow for site-specific labeling with FAM-maleimide (Lumiprobe Life science solutions). MBP-CTD fusion proteins were naturally devoid of cysteines, and a single cysteine residue at position 51 in full length SARS-CoV-2 Nsp1 was mutated to valine (SARS-CoV-2-Nsp1_C-51-V). This mutation had no effect on the ability of SARS-CoV-2 Nsp1 to inhibit mRNA translation in rabbit reticulocyte lysates, indicating that the protein is functional (data not shown). 100 µl protein at 50 µM was buffer exchanged into 40 mM HEPES-KOH (pH 7.5), 120mM KOAc, 5mM Mg(OAc)_2_, and a 10x molar access of FAM- maleimide and equal concentration of TCEP was added and the reaction mix incubated at RT for 2-3 hours protected from light. The reaction was desalted to remove free dye into buffer containing 40 mM Hepes-KOH (pH 7.5), 120 mM KOAc, 5 mM Mg(OAc)_2_, 2 mM DTT. Final protein concentrations and labeling efficiencies were calculated by absorption at 280 nm and 495 nm. Labeling efficiencies were generally > 80%.

### Ribosome purification

Mammalian 40S ribosomal subunits were purified from rabbit reticulocyte lysate (RRL) (Green Hectares). Briefly, RRL was thawed on ice, supplemented with 1 tablet of cOmplete EDTA-free protease inhibitor and ribosomes were pelleted through a 1 M sucrose cushion (in 50 mM KCl, 3 mM MgCl2, 2 mM DTT, 20 mM Tris-HCl pH7.5) over night at 40,000 rpm and 4 °C. Pellets were resuspended in 1 ml Buffer A (50 mM KCl, 3 mM MgCl2, 2 mM DTT, 20 mM Tris-HCl (pH7.5)) supplemented with proteinase inhibitor tablet and 10 µl RNasin (Promega). Fresh puromycin was added to a final concentration of 1 mM, followed by incubation for 20 mins on ice, then 15 mins at 37 °C. KCl was added to a final concentration of 0.5 M, and the ribosomes were layered onto a 10-30% continuous sucrose gradient in Buffer B (0.5M KCl, 3 mM MgCl2, 2 mM DTT, 20 mM Tris-HCl pH7.5). Samples were separated by centrifugation for 17 hrs at 22,000 rpm and 4 °C in an SW28 rotor, followed by fractionation using a BioComp Fractionator. Fractions containing 40S peak were pooled and ribosomal subunits pelleted through a 1 M sucrose cushion (in 110 mM KCl, 3 mM Mg(OAc)_2_, 20 mM Tris-HCl (pH 7.5)) over night at 40,000 rpm and 4 °C in a Ti50.2 rotor. Ribosome pellets were resuspended in 110 mM KCl, 3 mM Mg(OAc)_2_, 20 mM Tris-HCl (pH 7.5), 0.25M sucrose. Concentration were determined by A260 (40S preps generally yielded concentrations of 3-4 µM) and ribosomes were aliquoted and stored at −80 °C.

### Fluorescence polarization assay

Binding experiments with FAM-labeled proteins were performed using the Biotek Synergy Neo2 multi-mode microplate reader. The final concentration of labeled protein was limiting (5 nM) and the concentration of ribosomes was varied between 0.05 and 1000 nM. Binding reactions were performed in 20 mM Hepes-KOH (pH 7.5), 120 mM KOAc, 5 mM Mg(OAc)_2_, 0.2 mg/mL BSA, 2% sucrose. Reactions were incubated for 2 hours at 30 °C to reach equilibrium before fluorescence polarization was measured (multiple later measurements showed that the equilibrium had been reached). Total fluorescence was measured to account for quantum yield effects. The fraction bound was calculated (incorporating quantum yield changes between free and bound fluorophore) and binding fitted to a quadratic equation, assuming a tight binding regime^61^. K_D_ values reported are averages -+ SD.

### *In vitro* translation assay

*In vitro* translation assays were performed using nuclease-treated rabbit reticulocyte lysate (Promega). Reactions were in 10 µl containing 2 µl RRL, 1 µM amino acids, 25 mM Hepes-KOH (pH 7.5), 125 mM KOAc, 1 mM Mg(OAc)_2_, and the indicated protein concentrations diluted in protein storage buffer. Reactions were incubated for 20 mins on ice, and mRNA translation started by adding 50 ng FLuc mRNA (Promega), followed by incubation at 30 °C for 2 hours. Luciferase activity was measured using Luciferase Assay reagent (Promega) and analyzed on a Biotek Synergy Neo2 plate reader.

### RNA extraction and Northern blot analysis

RNA was extracted from HEK293T cells using Trizol Reagent (Invitrogen) according to the manufacturer’s instructions. 1 μg of total RNA was resuspended in 2x formamide RNA loading buffer, heated for 3 mins at 90C and separated on a 6% denaturing PAGE gel (Invitrogen). The RNA was then transferred to a HyBond-N+ nylon membrane (GE Life Sciences) using an electrophoretic transfer apparatus (Idea Scientific), UV-crosslinked at 120 mJ and blocked for 2 hours rotating at 42°C using ULTRAhyb Oligo hybridization and blocking buffer (Thermo Fisher). Blots were probed rotating overnight at 42°C with ^32^P-labeled DNA probes prepared as described below, and washed four times in 2x saline-sodium citrate (SSC) buffer with 0.5% SDS for 10 minutes at 42°C. The blots were imaged using a phosphor screen and Typhoon 9400 scanner (GE Life Sciences). DNA oligos used as Northern probes were ordered from IDT (GFP: 5’-CCG GAC ACG CTG AAC TTG TGG CCG TTT-3’, S6: 5’-TAT GGA ACG CTT CAC GAA TTT GCG TGT CAT CC-3’). 100 pmol of DNA was incubated with 2 μL 5 mCi [γ32-P]ATP (PerkinElmer) and 4 μL T4 PNK (New England BioLabs) in a 100 μL reaction for 2 hours at 37°C and purified using Micro-Bio P-30 spin columns (Bio-Rad). The radiolabeled probe was heated to 90°C for 2 minutes and resuspended in 10 mL of ULTRAhyb Oligo hybridization buffer (Thermo Fisher). Blots were incubated with 10 mL of the resuspended probe overnight at 42°C.

### NMR

To express ^15^N-labeled protein, M9 minimal media was used inoculated with ^15^N-ammonium chloride. The protein was expressed and purified as described above and dialyzed in NMR buffer (25 mM NaP, 250 mM NaCl, 2 mM DTT, 0.02% w/v NaN3, pH 7.0). ^15^N R_1_ and R_2_ relaxation rate constants of Nsp1 were measured on a triple-resonance BRUKER 600MHz NMR cryo-probe spectrometer at 25 °C. The relaxation sampling time points were 80, 100, 200, 300, 400, 500, 600, 700, 800, 900, and 1000 ms for R_1_ and 16, 33, 67, 135, 169, 203, 237, and 271 ms for R_2_. Data processing was performed using the NmrPipe^68^ package. Data fitting was done using GraphPad PRISM (V. 9).

### Comparative homology modeling for Nsp1-ribosome complexes

We used RosettaCM^57^ to model the SARS-CoV, MERS-CoV, HuCoV-HKU1, HuCoV-OC43 and MHV Nsp1-ribosome complexes. Detailed computational methods, PDB files of all homology models, and all associated analysis scripts are available at https://github.com/glasgowlab/Interface_Energy_nsp1. Briefly, input structures to serve as templates for RosettaCM were prepared by running Rosetta relax^69^ with backbone constraints in a 150 Å radius around the Nsp1 C-terminal peptide on eight experimentally-solved SARS-CoV-2 structures (PDB IDs 6ZLW, 6ZM7, 6ZME, 6ZMI, 6ZMO, 6ZMT, 6ZN5, 6ZOJ)^12^. The five viral Nsp1 C-terminal sequences were partial threaded on the eight templates, then 1500-2000 homology models were hybridized per Nsp1 sequence using RosettaScripts^70^, Nsp1 fasta files, and the threaded templates as inputs. The models were ranked according to total score by the Rosetta energy function. The four models with the lowest total energies for each Nsp1 sequence were then structurally aligned^71^ to the experimentally-solved Nsp1 C-terminal peptide fragment in the SARS-CoV-2 Nsp1-ribosome complex (PDB ID 6ZLW), and 20 total Nsp1-ribosome models were relaxed and scored. The lowest-energy complex models were then analyzed using PyRosetta to characterize the predicted Nsp1-ribosome interactions. Each nsp1-ribosome complex model was scored using the Rosetta energy function with the *rna_res_level_energy4* weights settings, and intermolecular interface energies and individual score terms were calculated for Nsp1, ribosomal proteins, ribosomal RNAs, and their interactions^72–74^.

## Acknowledgements

We thank members of the Steckelberg lab for providing valuable feedback. This work was supported by NIH grants R21AI171827 (A.-L.S. and M.A.H.) R00GM135529 (A.G.), R35GM118070 (J.S.K.) and R01AI133348 (J.S.K.).

