## Supplement for "An evolutionarily conserved strategy for ribosome binding and inhibition by β-coronavirus non-structural protein 1"

**Supplementary Material**

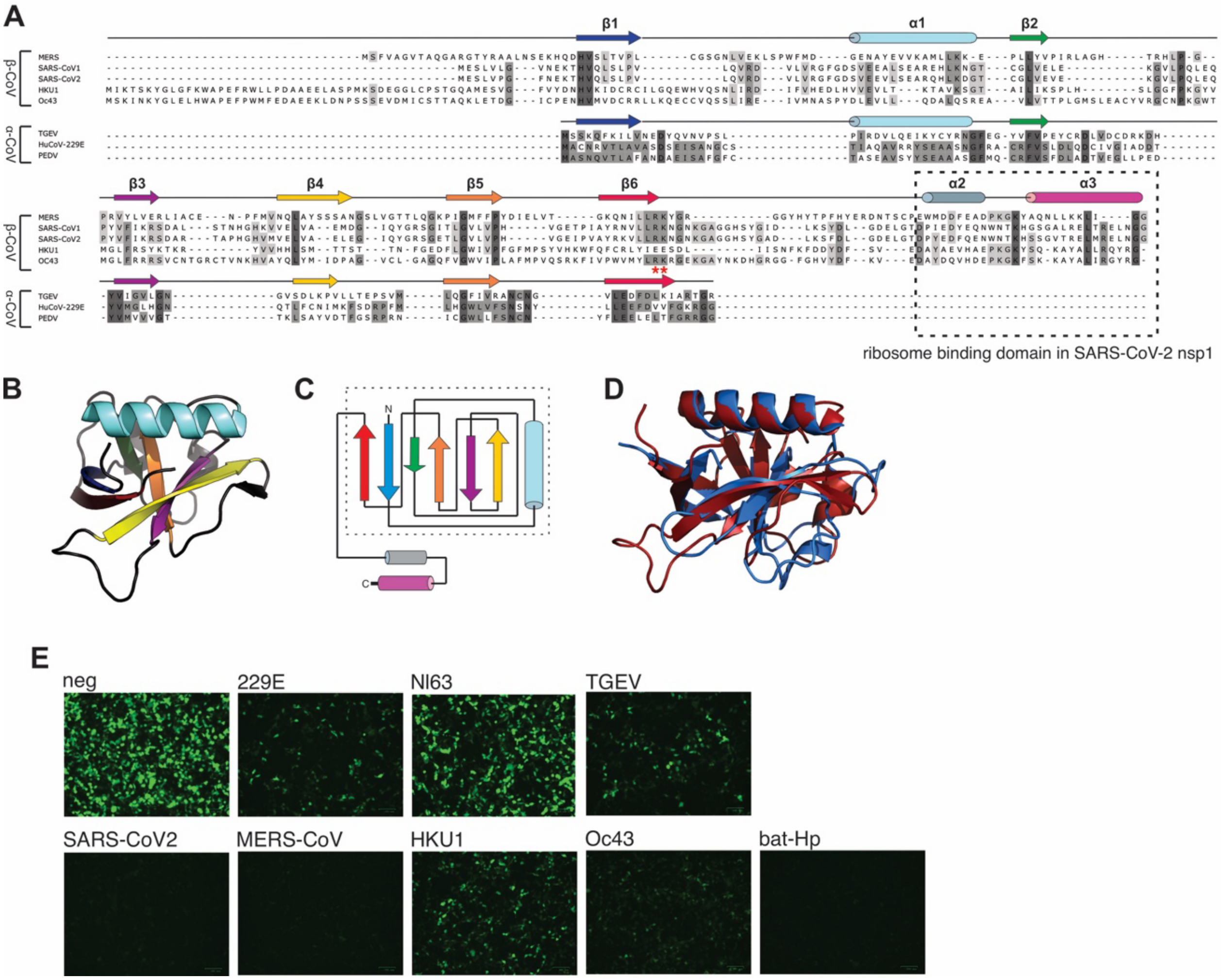


**Figure S1: Sequence diversity but structural and functional conservation of Nsp1 from α- and β-coronaviruses.** A) Sequence alignment of Nsp1 from various α- and β-coronaviruses. Multiple sequence alignments of α-CoV Nsp1 and β-CoV Nsp1 sequences were performed independently, and manually aligned based on structure conservation. B) N-terminal globular domain of SARS-CoV-2 (pdb: 7k7p) C) Schematic representation of the SARS-CoV-2 NTD fold, secondary structure elements are colored to match B. Box depicts NTD, which is structured in solution D) Overlay of the NTD of SARS-CoV-2 Nsp1 (in red, pdb 7k7p) and full-length TGEV Nsp1 (in blue, pdb: 6IVC). Note the similarity in 3D structure despite considerable sequence variation. E) Host shutoff in HEK293T cells transiently expressing Nsp1 from the indicated viruses and EGFP reporter gene.

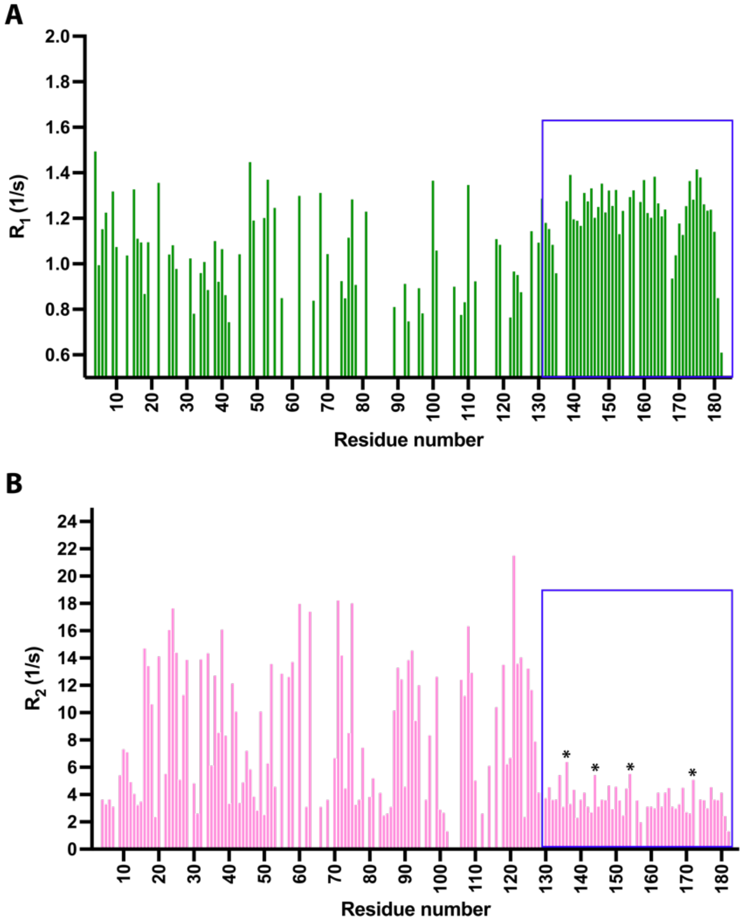


**Figure S2: NMR relaxation experiments show the intrinsically disordered CTD of Nsp1**. A) $^{15}$N longitudinal relaxation rates $R_1$. B) $^{15}$N transverse relaxation rates $R_2$. Analysis of $R_2$ relaxation data showed an average $R_2$ value of 6.8 $s^{-1}$ and a decrease in relaxation rates from residues G-127. Measurements were conducted at 600 MHz field strength. The magenta box marks the CTD. The asterisks denote CTD residues His-134, Ser-141, Asp-152, Thr-170, that have higher than average $R_2$, demonstrating structural compaction and/or exchange on the μs-ms timescale.

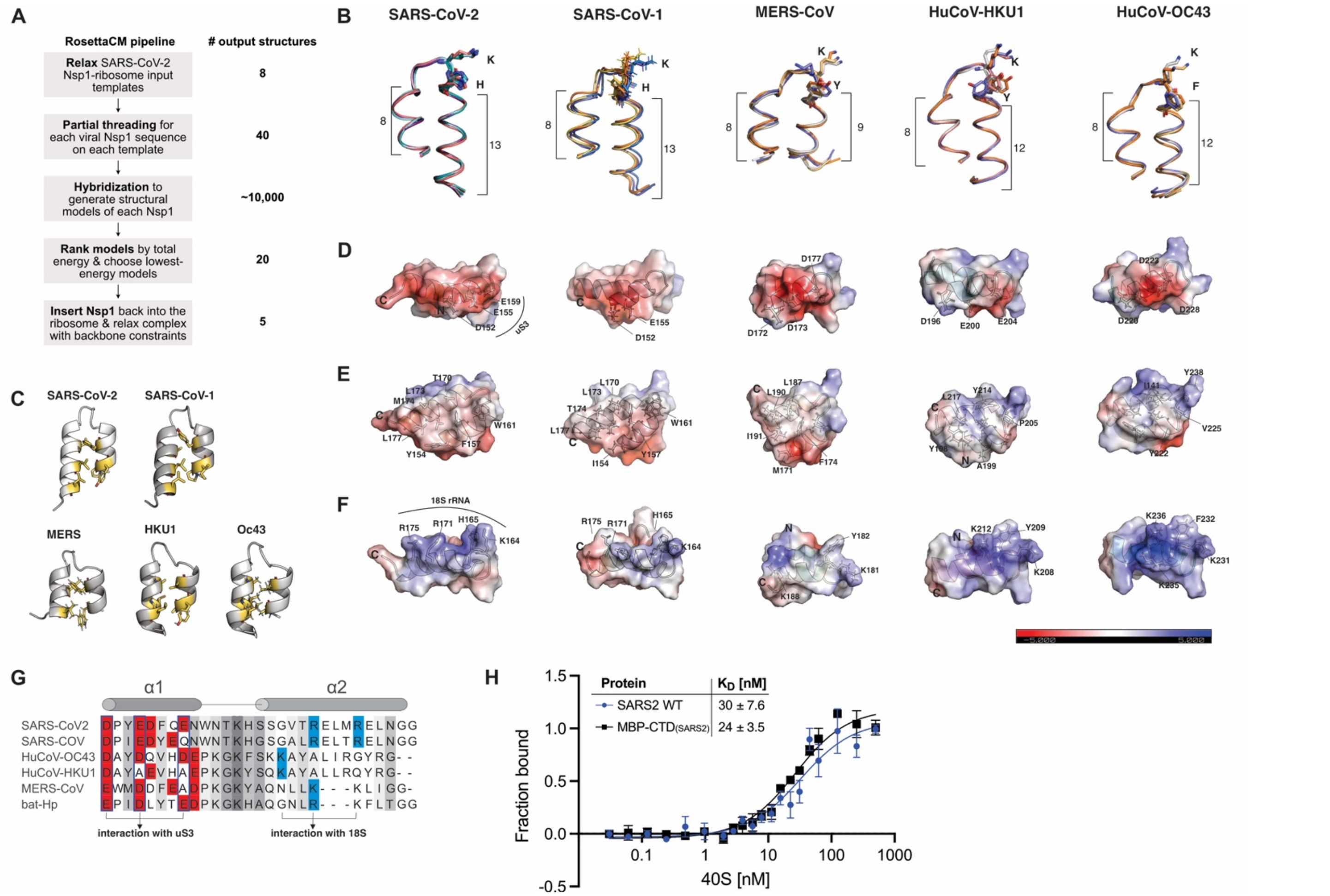


**Figure S3: Modeling of the C-terminal domain from divers β-CoV Nsp1.** A) Overview of the homology modeling pipeline. B) Overlay of the Rosetta models of the Nsp1 CTD from the indicated β-CoVs. The conserved KH or KY/F motif is shown in the loop. Numbers indicate the length (in amino acids) of α-helix 1 and 2. C) Hydrophobic residues at the helix interface stabilize the Nsp1 CTD. D-F) Electrostatic surface representation of the Nsp1 CTD models shows partial surface charge conservation between divers Nsp1 variants. Electrostatics were calculated using the Adaptive Poisson-Boltzmann Solver (APBS) in pymol. The surface colors are fixed at red (-5) and blue (+5). G) Sequence alignment of the Nsp1 CTD from SARS-CoV, SARS-CoV-2, MERS-CoV, HuCoV-HKU1, HuCoV-OC43 and bat-Hp nsp1. Negatively charged amino acids in helix 1 are depicted in red, and positively charged amino acids in helix 2 are depicted in blue. Note that the charged amino acids are positioned differently along the helices. H) Equilibrium binding measurements of FAM-labeled SARS-CoV-2 Nsp1 and MBP-$CTD_{SARS2}$ to purified rabbit 40S ribosomal subunits in buffer containing 250 mM $K^+$. n=3, error bars=SEM.

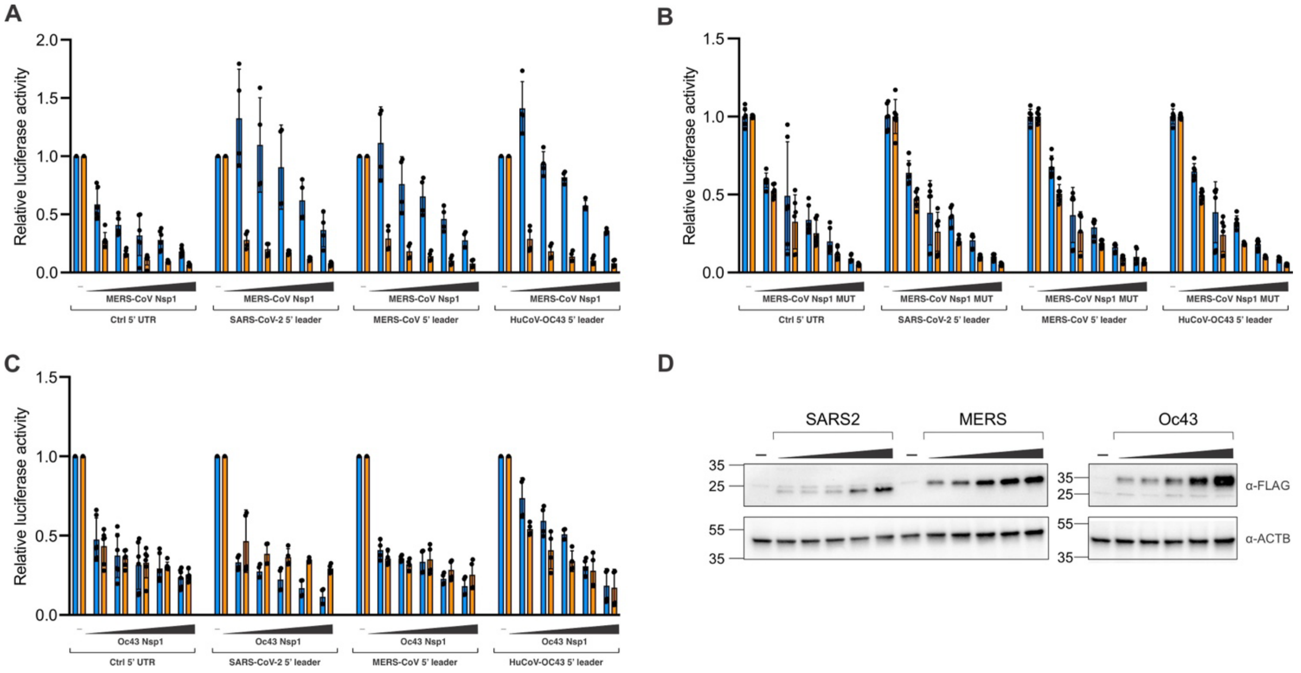


**Figure S4: No virus-specific RNA-protein interaction modulates MERS-CoV and HuCoV-OC43 Nsp1 function.** A-C) Nsp1-dependent host shutoff and translation boost in HEK293T cells transiently expressing MERS-CoV Nsp1 (A), MERS-CoV Nsp1 (RK-146/7-AA) (B), or HuCoV-OC43 Nsp1 (C) and FLuc reporters with the indicated 5' UTRs. NLuc was co-transfected as an internal control. N=6, error bars = SD. D) Representative immunoblot analysis showing the expression levels of Nsp1.
